# Efficacy of a phage cocktail preparation for chronic lung infection with *Pseudomonas aeruginosa* in mice

**DOI:** 10.1101/2023.04.25.538306

**Authors:** Ru-yue Gao, Xin Tan, Yong-jun Pan, Jia-lin Yu, Ying-fei Ma

## Abstract

*Pseudomonas aeruginosa* is a leading cause of hospital-acquired infections, and the emergence of multi-drug resistant strains has prompted the search for alternative treatments such as phage therapy. In this study, we combined host range and genomic information to design a four-phage cocktail that effectively killed several clinical strains (79%, 23/29) of *P. aeruginosa*. We demonstrated that the cocktail, composed of three novel phages (PA_ZH1, PA_GL1, and PA_CQ9) and one previously characterized phage (PA_LZ7), was able to lyse *P. aeruginosa* both in planktonic cultures and in alginate microbeads (an in vivo-like biofilm model). Additionally, we showed that the phage cocktail administered intranasally or intraperitoneally effectively rescued mice from chronic lung infection with *P. aeruginosa*. Our work explores the potential use of phages as an alternative therapeutic agent against chronic lung infections caused by *P. aeruginosa* strains.

## Introduction

*Pseudomonas aeruginosa* infections are a severe concern in healthcare settings, especially for patients with compromised immune systems or those who require prolonged hospital stays^1^. The ability of *P. aeruginosa* to form biofilms in the lungs of patients such as cystic fibrosis (CF) is particularly concerning, as this can lead to chronic infections that are difficult to eradicate^2^. The extracellular matrix of the biofilm can prevent antibiotics from reaching the bacteria within, and the bacteria themselves may be less susceptible to antibiotics due to the protective nature of the biofilm^3^. Additionally, bacteria within a biofilm may exhibit altered gene expression and metabolism, further complicating treatment strategies^2, 3^. Furthermore, the emergence of multi-drug resistant (MDR) strains of *P. aeruginosa* adds another layer of difficulty in treating infections caused by this pathogen^1^. Thus, there is an urgent need for the implementation of novel antimicrobial agents to treat *P. aeruginosa* infections.

The therapeutic potential of phages for bacterial infections has been highlighted in numerous recent studies^4–6^. Phages have many advantages over traditional antibiotics. One advantage is their host specificity, which means that they only target specific bacterial strains, leaving the rest of the microbiota undisturbed^7^. Additionally, phages can self-replicate in the presence of host bacteria, allowing for a higher concentration of phages to accumulate and ultimately kill the bacteria^7^. Another critical advantage of phage therapy is that it is not limited by antibiotic resistance^4, 7^. This is due to the fact that phages bind specific receptors on the bacterial cell surface, and bacteria must mutate those receptors to become resistant^8^. However, even if one strain of bacteria evolves resistance to a specific phage, a phage cocktail containing multiple phages can effectively overcome this issue^5^. Moreover, phages have been shown to degrade biofilms^9, 10^. Despite these advantages, most studies on phage therapy have focused on acute infections, such as lung, bloodstream and skin, and soft tissue infections^11^. However, few studies have explored the application of phages in chronic lung infections. Therefore, there is a need to investigate the potential of phage therapy for the treatment of chronic pneumonia. To address this knowledge gap, we chose to investigate the efficacy of phage therapy in a mouse model of chronic lung infection caused by *P. aeruginosa*.

Our study aims to determine the efficacy of phage cocktail therapy in the treatment of chronic lung infection with *P. aeruginosa* in mice. In addition, the efficacy between the intranasal (i.n.) and intraperitoneal (i.p.) delivery routes were compared to determine whether different delivery methods have different efficacy.

## Result

### Biological characteristics and lytic effect of isolated phages

We isolated three phages using the PAO1 strain as the host: PA_ZH1, PA_GL1, and PA_CQ9 from sewage water samples. The phages produced clear and round plaques with diameters ranging from 1.3 to 3.6 mm, and the largest clear plaque was formed by PA_GL1 (**Figure 1A**).

**Figure 1.**
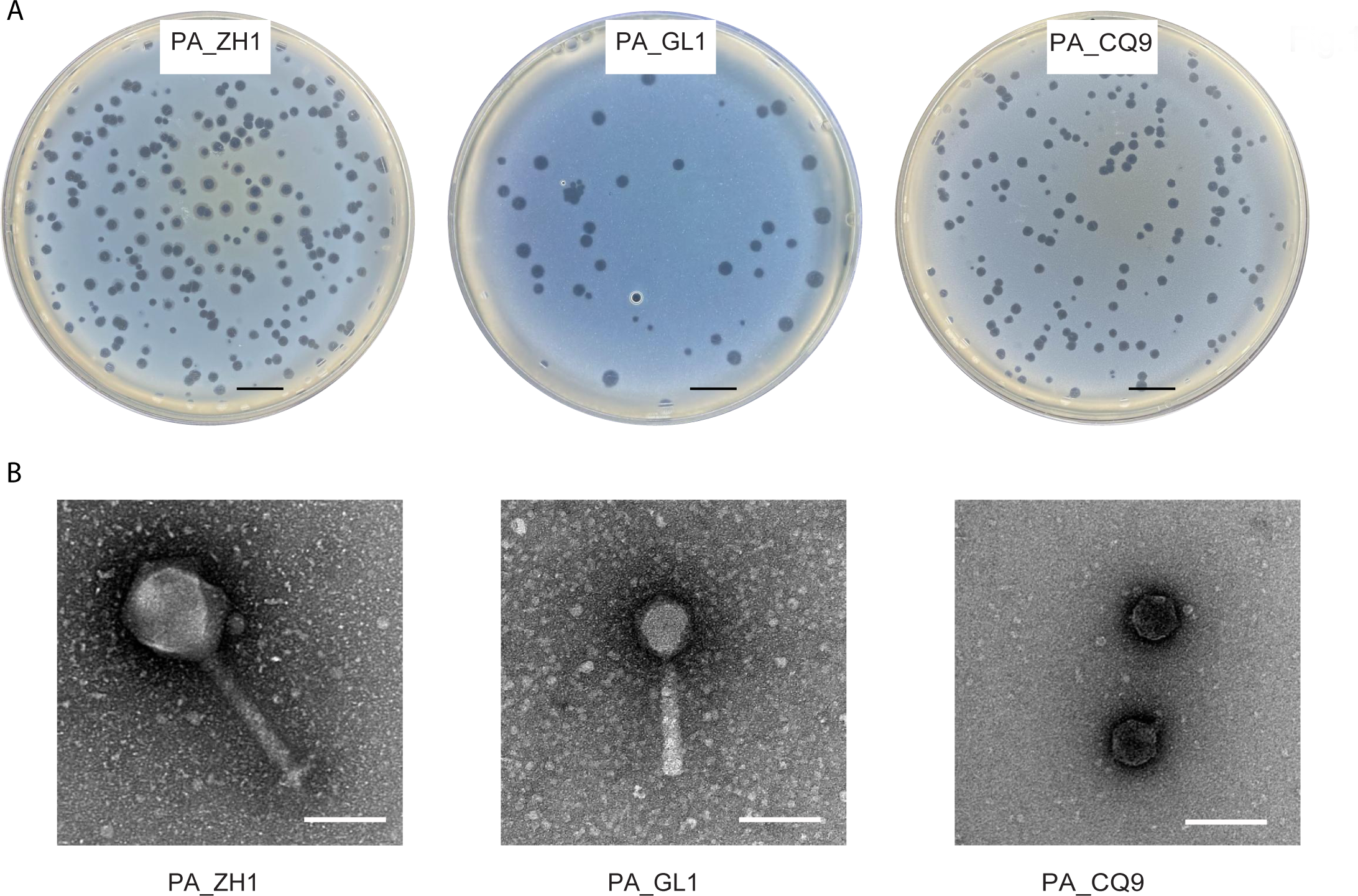
Plaque and transmission electron microscopy (TEM) morphology of phages. (A) Plaque morphologies of phages PA_ZH1, PA_GL1 and PA_CQ9. Scale bars represent 1 cm. (B) TEM morphologies of phages PA_ZH1, PA_GL1 and PA_CQ9. The phages were stained with 2% phosphotungstic acid and visualized with TEM. Scale bars represent 100 nm.

Transmission electron microscopy (TEM) images of these phages are shown in **Figure 1B**. All belong to the order of *Caudovirales*. PA_ZH1 belongs to myovirus, and possesses an icosahedral head of 110 nm in diameter and a tail 175 nm long; PA_GL1 are myovirus with a regular icosahedral head of 75 nm in diameter and a tail of 140 nm; PA_CQ9 belongs to podovirus, and possesses an icosahedral head of 55 nm and a tail of 13 nm.

### Genome characterization and analysis of the selected phages

The genomic sequences of phage PA_ZH1, PA_GL1, and PA_CQ9 have been deposited in GenBank with accession numbers OQ828465, OQ828463, and OQ828464, respectively. Phage PA_ZH1 has a genome length of 278,577-bp and is affiliated with the *Phikzvirus* genus. Its nucleotide sequence shows 99% identity with *Pseudomonas* phage phiKZ (GenBank: NC_004629) and KTN4 (GenBank: KU521356). Phage PA_GL1 has a genome length of 65,283-bp and is affiliated with the *Pbunavirus* genus. Its nucleotide sequence shows around 97% identity with *Pseudomonas* phage PB10 (GenBank: OP831166) and NH-4 (GenBank: NC_019451). Phage PA_CQ9 has a genome length of 45,672-bp and is affiliated with the *Bruynoghevirus* genus. Its nucleotide sequence shows around 98% identity with *Pseudomonas* phage Pa222 (GenBank: MK837011) and PSA34 (GenBank: MZ089739), which is a LUZ24-like phage.

The linear genetic map of PA_ZH1, PA_GL1, and PA_CQ9 can be seen in **Figure 2**. Analysis of the phage whole genome shows that they possess a series of genes encoding common phage-related features, including DNA polymerase, DNA helicase, tail and head structure proteins. They also contain genes encoding the host lysis protein, endolysin and holin. Notably, the *in-silico* analysis did not reveal any putative virulence or antibiotic resistance or integrase sequences in the genome of these phages.

**Figure 2.**
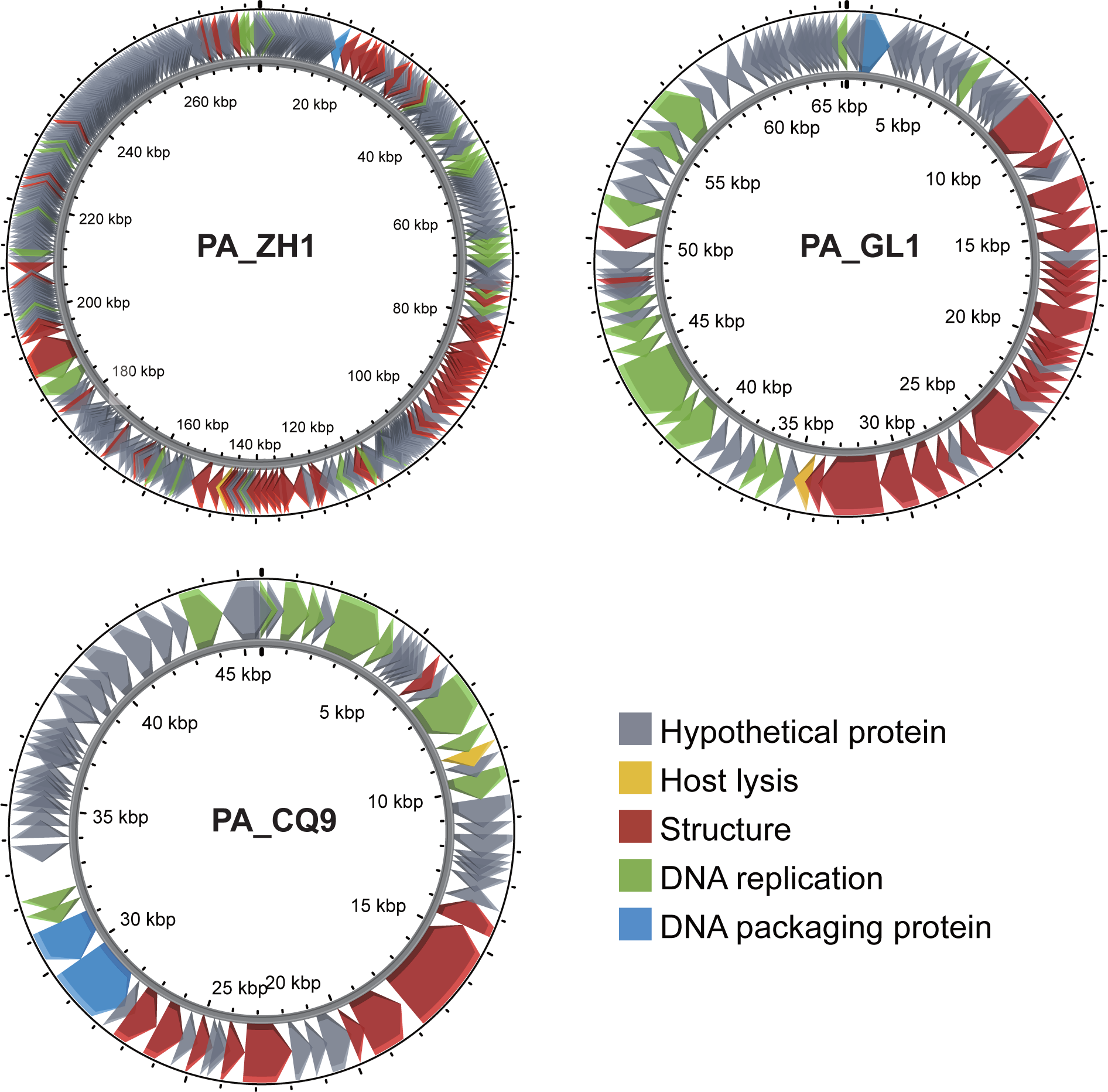
Genome annotation of *Pseudomonas* phages PA_ZH1, PA_GL1 and PA_CQ9. The color of the ORFs refers to five modules: phage structure, red; host lysis, yellow; DNA packaging, blue; DNA replication, green; and hypothetical proteins, grey.

### Definition of a genetically diverse phage cocktail

To compose a phage cocktail that displays a broad host range and genetic diversity, we selected four virulent phages, PA_ZH1, PA_LZ7, PA_GL1, and PA_CQ9. PA_LZ7 is previously characterized *P. aeruginosa* virulent phages isolated in China^12^ that are 66, 155-bp, and that display 97% identity over 99% of its genome length with PA_GL1. Here, we refer to the mixture of all four phages as the “phage cocktail.”

We determined the host range of the four phages against a panel of 17 clinical *P. aeruginosa* strains (**Table 1**). As expected, each phage had a distinct host range, with no individual phage being able to lyse all strains in the aforementioned collection. However, the phage cocktail showed the ability to lyse 79.3 % (23/29) of the tested *P. aeruginosa* strains (**Table 1**).

**Table 1.** Phage lytic spectra on the bacterial strains used in this study. Five microliters of the indicated single phage or phage cocktail was spotted on a lawn of each specific bacterial host; the plates were observed after overnight incubation at 37°C. +, clear plaque; -, no plaque; T, turbid plaques.

| Bacterial Strain | Infectivity |  |  |  |  |
| --- | --- | --- | --- | --- | --- |
|  | PA_ZH1 | PA_LZ7 | PA_GL1 | PA_CQ9 | Cocktail |
| PAO1 | + | + | + | + | + |
| RPA01B | + | T | + | + | + |
| RPA02B | T | + | + | T | + |
| RPA03B | + | T | T | T | + |
| RPA04B | + | - | T | + | + |
| 53776 | T | + | - | + | + |
| PA01B | + | - | - | T | + |
| PA02B | - | + | + | - | + |
| PA03B | + | - | - | + | + |
| PA04B | + | + | - | T | + |
| PA05B | + | + | T | T | + |
| PA06B | + | - | - | + | + |
| PA07B | T | T | T | - | + |
| PA08B | - | + | + | T | + |
| PA09B | - | + | + | T | + |
| PA10B | - | + | T | + | + |
| PA11B | + | + | - | + | + |
| PA12B | + | + | - | - | + |
| PA13B | + | + | + | - | + |
| PA14B | + | + | - | - | + |
| PA15B | T | T | T | - | T |
| PA16B | + | + | + | + | + |
| PA17B | - | - | - | - | - |
| PA18B | - | - | T | - | T |
| PA19B | - | - | - | - | - |
| PA20B | + | T | - | + | + |
| PA21B | - | - | T | T | T |
| PA22B | - | T | T | - | T |
| PA23B | + | + | T | T | + |

### In vitro characterization of the cocktail

The lysis kinetics of *P. aeruginosa*strain PAO1 cultures infected with each phage and cocktail was followed by monitoring the optical density (OD) at 600 nm (OD_600_) over time (**Figure 3A**). At a multiplicity of infection (MOI) of 1:100, all phages caused a decrease in the OD_600_ at 1 h post-infection. After 8 h of incubation, the OD_600_ of single phage groups tended to increase, due to the growth of resistant bacteria. However, in the cocktail group, the potent inhibition effect was still observed after 24 h co-incubation, indicating that the cocktail can efficiently prevent the outgrowth of phage-resistant variants.

**Figure 3.**
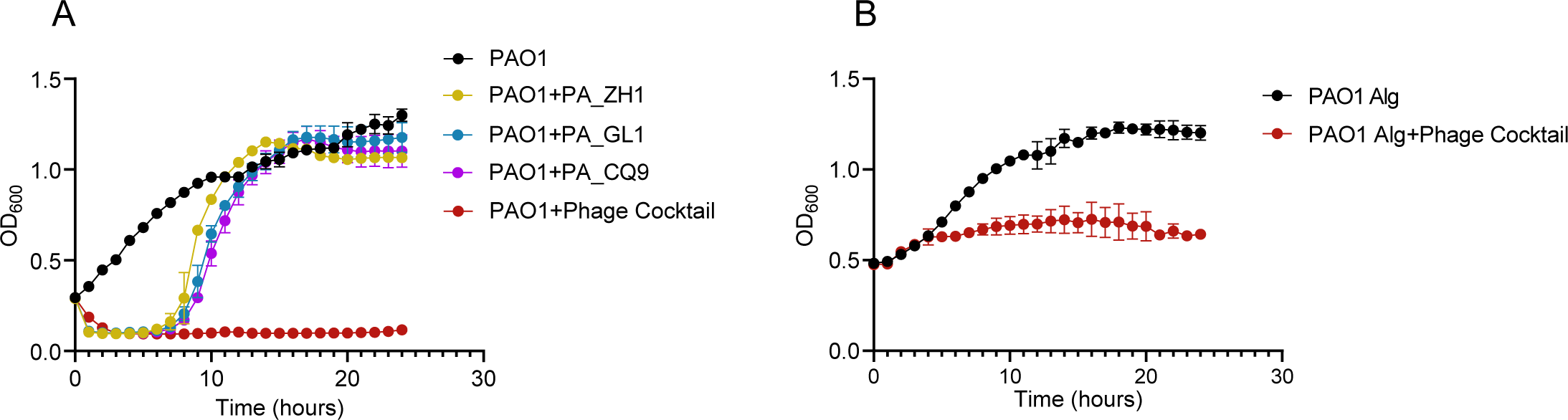
Lytic effect of individual phage and phage cocktail against PAO1 in vitro. (A) Growth kinetics of planktonic PAO1 in presence of phages. Exponentially growing PAO1 (OD600=0.3) were infected by the indicated phages or phage cocktail, each at an MOI of 1. (B) Growth kinetics of PAO1 in alginate beads in presence of phage cocktail. Data represent the mean with standard deviation. The results of one out of three independent experiments are shown.

The capability of the phage cocktail to inhibit bacterial growth in an in vivo-like biofilm model, where *P. aeruginosa* was encapsulated in alginate beads (**Figure S1**), was also tested (**Figure 3B**). The phage cocktail caused a decrease in OD_600_ at 5 h post-infection, and a strong inhibitory effect was still observed after 24 h of co-incubation, indicating the cocktail’s efficiency in inhibiting bacterial growth in biofilm.

### Therapeutic efficacy of phage cocktail in a mouse model of chronic *P. aeruginosa* lung infection

To mimic a chronic infection that biofilm is typically formed in the lungs of patients, mice received intratracheal inoculations with the *P. aeruginosa* PAO1 strain embedded in alginate beads^2, 13^.

To evaluate the effect of the phage cocktail against bacterial infection, administration via the i.n. or i.p. route with the phage cocktail was started 30 min after infection and was repeated daily for seven administrations. The phage cocktail was administrated as follows: 5*10^7^ PFU for each phage, with a total of 2*10^8^ PFU per dose (See details for phage preparations in **Table S1**).

In the non-phage-treated control group, the mice showed a significant decrease in activity, reduced food and water intake and weight loss, presence of unruly hair, arched back and chills. In contrast, the general condition of the mice in the phage-treated groups was better than that of the non-phage-treated control groups. The survival rate of mice in the i.n. phage-treated group was significantly higher than that of mice in the non-phage-treated control group (*P*<0.01) (**Figure 4A**). Similarly, the survival rate of mice in the i.p. phage-treated group was considerably higher than that of mice in the non-phage-treated control group (*P*>0.05) (**Figure 4B**).

**Figure 4.**
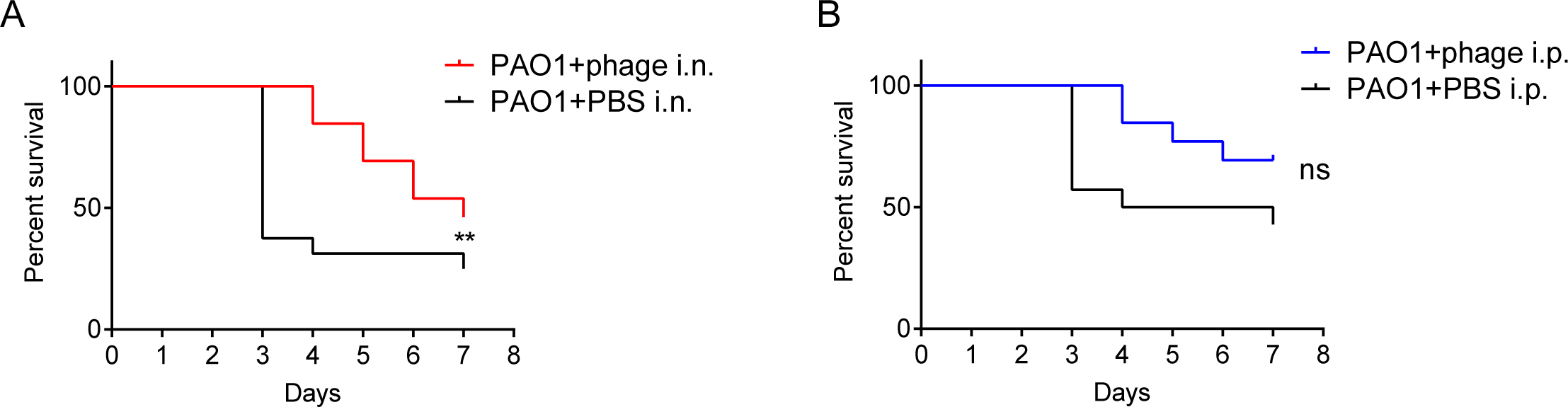
Efficacy of phage cocktail in a mouse model of chronic *Pseudomonas aeruginosa* lung infection. Survival rate was evaluated after intranasal (i.n.) or intraperitoneal (i.p.) administration of phage cocktail (phage treatment groups, n=13; PAO1 i.n. group, n=16; PAO1 i.p. group, n=14). Phages treated with a dose of 2*10^8^ PFU (MOI = 28) at 30 min post-infection (*, *P*<0.05; **, *P*<0.01).

### Bacteriological burden, histological changes and cytokine analysis of lung tissues

Mice in the non-phage-treated groups exhibited significantly higher bacterial loads, reaching approximately 10^6^ CFU/lung, while the bacteria burden in phage-treated groups was almost decreased to 0 CFU/lung (*P*<0.01) (**Figure 5**).

**Figure 5.**
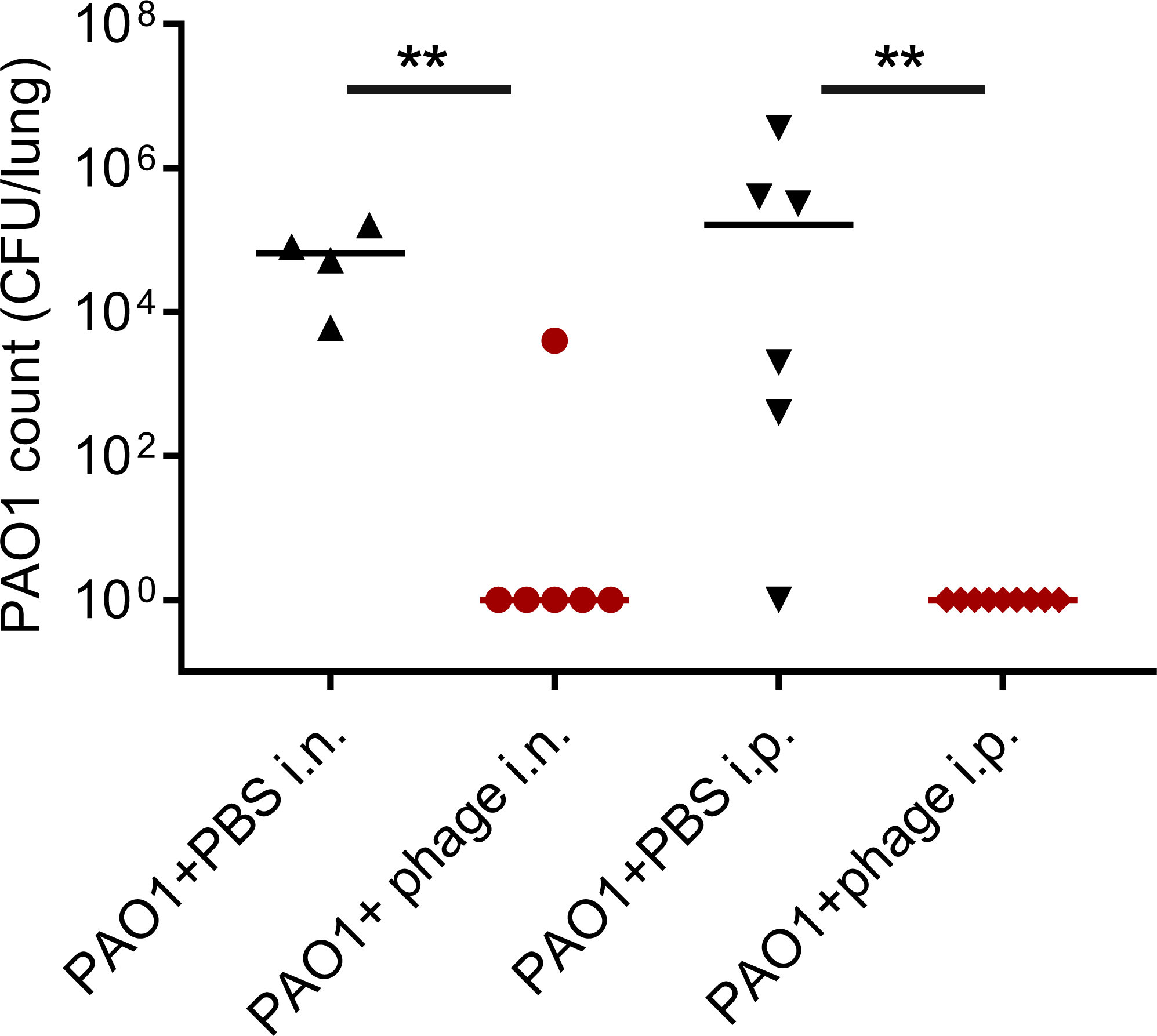
Bacterial burden in the lung 7 days post-infection. Mice in the bacteria-infected group and phage-treated group were sacrificed 7 days post-infection and the bacterial burden in lungs was measured by using bacterial colony counting. Individual data and the medians were shown in the figure. The comparation was performed with Mann-Whitney test (**, *P*<0.01).

Lung tissue from the mice of non-phage-treated groups showed widened alveolar septum, fibrosis and a large number of inflammatory cells, with hemorrhage and red blood cell infiltration in the alveolar cavity (**Figure 6G and 6H**). In contrast, lung tissue from the mice of the i.n. phage-treated group showed typical lung structure (**Figure 6E**). Lung tissue from mice of the i.p. phage-treated group showed a little inflammatory cell exudation, but the level of inflammation was less than that of the non-phage-treated group, and the fine bronchial structure was generally intact (**Figure 6F**).

**Figure 6.**
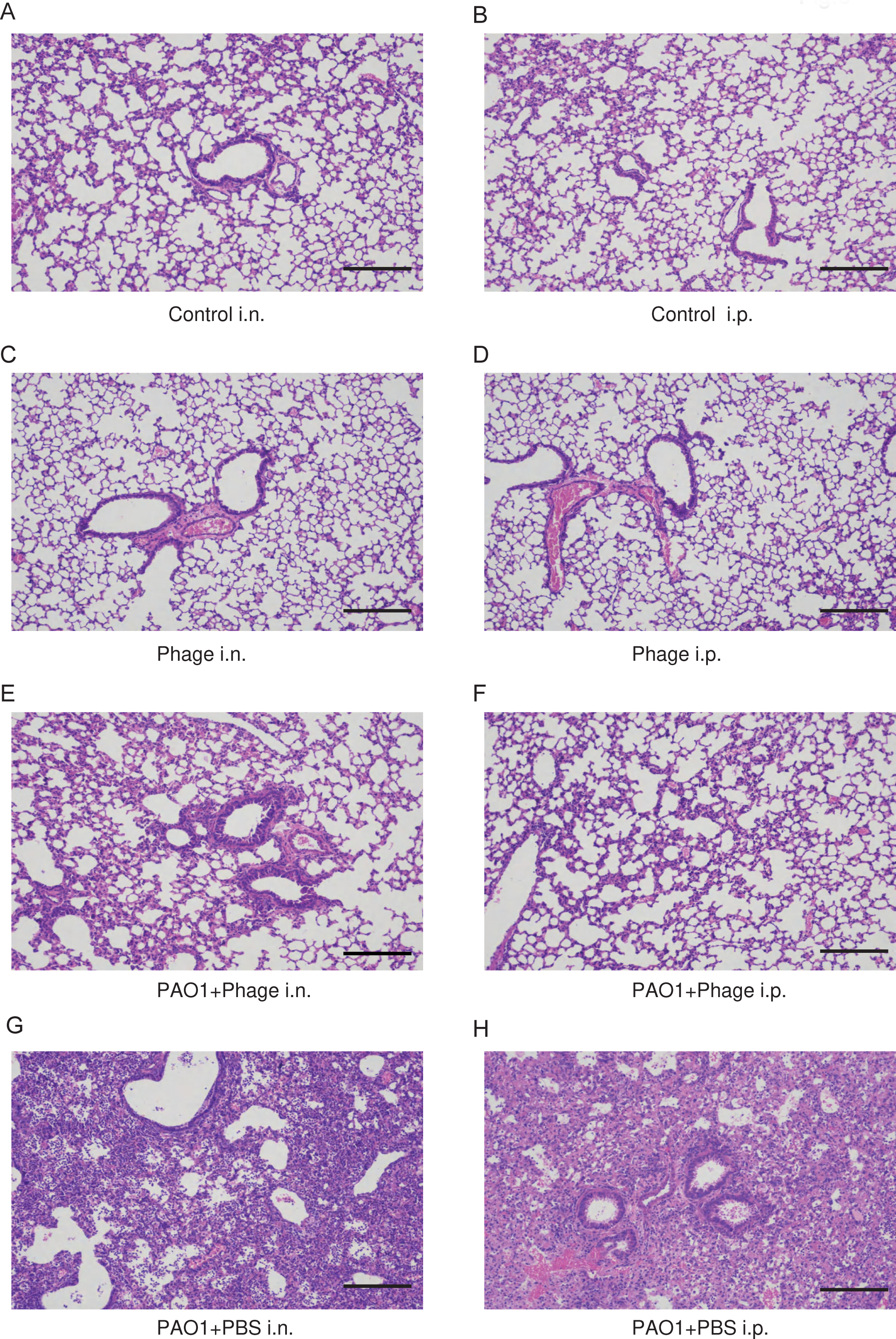
Histology analysis of mice lung tissue. (A) control i.n. group; (B) control i.p. group; (C) phage i.n. group; (D); phage i.p. group; (E) PAO1+PBS i.n. group; (F) PAO1+PBS i.p. group; (G) PAO1+phage i.n. group; (H); PAO1+phage i.p. group. Hematoxylin and Eosin (H&E) staining was used for histopathologic examination. Scale bars represent 200 μm.

Cytokines in the lung homogenate of the mice from different groups were measured on day 7 **(Figure 7**). The result showed that the levels of all cytokines were elevated in the lung homogenate of PAO1 infected mice, while i.p. phage treatment significantly (*P*<0.05) reduced the excessive TNF-α, IL-6, IL-10, IL-1β, G-CSF, CCL2/JE/MCP-1, CCL3/MCP-1α and CCL4/MIP-1β release caused by bacterial overgrowth **(Figure 7**). IFN-γ also showed decreased level (*P*>0.05). I.n. phage treatment also reduced (*P*>0.05) the production of these cytokines. The expression of cytokines remained at basal levels in the mice from both vehicle and phage control groups **(Figure 7**).

**Figure 7.**
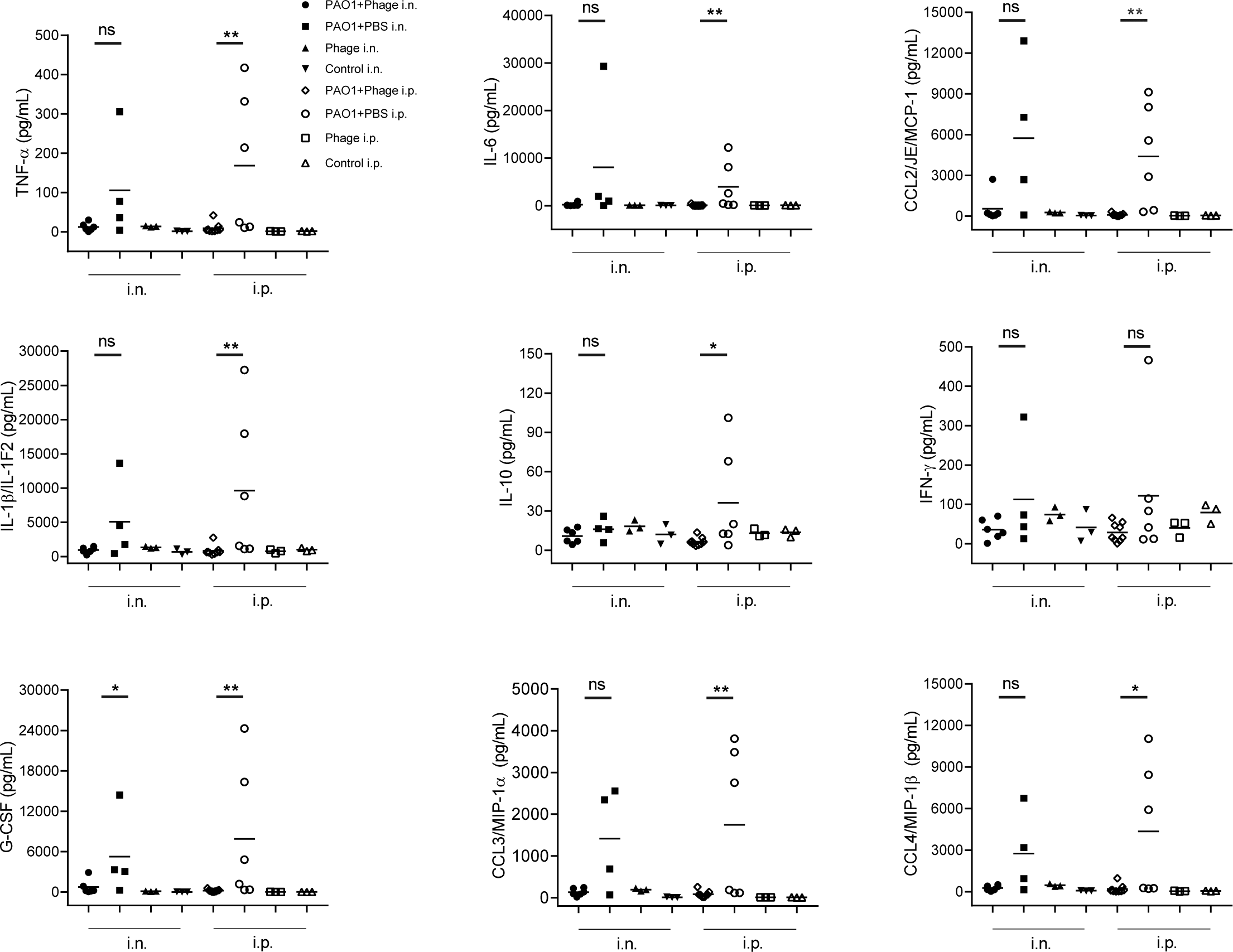
Cytokine levels in mice lung tissue homogenate 7 days post-infection. The lung tissue homogenates of the mice were collected 7 days post-infection. Cytokines concentrations were measured by Luminex multiplex cytokine assay. The medians were shown in the figure. The comparation was performed with Mann-Whitney test (*, *P* < 0.05; ** < 0.01).

## Discussion

*P. aeruginosa*, a common opportunistic pathogen, is now generally recognized as one of the most causative agents of hospital-acquired infections^1^. MDR *P. aeruginosa* has become more and more common in recent years, posing severe challenges to global health^1, 14^. Moreover, *P. aeruginosa* was included in the World Health Organization’s list of superbugs that urgently need novel antibiotics in 2017^1^. The search for novel treatments for MDR *P. aeruginosa* has become a hot topic. Recently, the therapeutic potential of phages for bacterial infections has been highlighted. Phage treatments of animals have been reported in many studies with positive outcomes^5, 11, 15^. In addition, personalized phage cocktails were successfully applied to treat MDR *P. aeruginosa* infections in clinical cases^6, 16–18^. Several controlled clinical trials also showed the safety and efficacy of phage therapy for burn wound infections, urinary tract infections and otitis^19–22^, these results suggest a promising future for phage therapy. However, there are limited reports on the application of phage in chronic lung infections.

In this study, we isolated and identified three phages, and analyzed phage morphology, genomic information, host range, antibacterial capacity in vitro. The host range of phage is usually narrow, which will impede their clinical efficacy. To tackle this problem, phage cocktails are applied to broaden the host range and make it more effective than mono-phage therapy^23–25^. Our study indicated that the combined infection spectrum of four phages was 72.2 % (13 of 18) against *P. aeruginosa* clinical isolates (**Table 1**), which represents a good coverage of *P. aeruginosa* strains.

Further, a cocktail of different types of phages that infect the host bacterium by different mechanisms is often used in practice, which can significantly reduce the chance of creating mutant strains that resist several phages at the same time. The receptors for phiKZ^26^, PB1-like phages^27, 28^, and PSA34 phage^29^ were identified as type IV pilin (T4P), lipopolysaccharide (LPS), and LPS, respectively. Since PA_ZH1 shared a high identity with phiKZ, PA_LZ7, and PA_GL1 shared a high identity with PB1, PA_CQ9 shared a high identity with PSA34, we hypothesized that the receptors for these phages might be T4P and LPS, respectively. The potent inhibition effect was observed after 36 h co-incubation with phage cocktail (**Figure 3A**), indicating that the cocktail can efficiently prevent the outgrowth of phage-resistant variants.

Biofilm is formed when *P. aeruginosa* in its planktonic state adheres to a solid surface, aggregates together and forms microcolonies, which form the primary structure of the biofilm by bonding to each other and then differentiating to form a mature biofilm^3^. *P. aeruginosa* progeny grown in the biofilm can switch into a state of low metabolism, low energy consumption and low activity, and therefore can escape detection by various immune cells and escape the action of antibiotics, thus generating antibiotic resistance ^2, 3^. The ability of *P. aeruginosa* to form biofilms in the lungs of patients such as CF is particularly concerning, as this can lead to chronic infections that are difficult to eradicate^2^. Recently, the alginate-based beads, a in vivo-like biofilm model, were successfully established to mimic the lung infections observed in CF patients^30, 31^. Treatment of *P. aeruginosa* in alginate beads with the phage cocktail demonstrated that phages could efficiently kill the bacteria within the biofilm (**Figure 3B**). Indeed, phages can disaggregate biofilms through different mechanisms, including replication within their host, leading to an increase in the number of phages and progressive removal of the biofilm, encoding depolymerases, which are responsible for the degradation of extracellular polymeric substances (EPS) matrix components and the penetration of phages into the biofilm^9, 32^. Interestingly, PA_ZH1 produced plaques with haloes, which are semi-transparent zones around the plaques (**Figure 1A**). This halo formation is regarded as a hallmark of depolymerase activity^33^, which can explain the anti-biofilm effect of the phage cocktail. However, no depolymerase coding genes were found in the genome of PA_ZH1 through bioinformatic analysis, further study is needed to investigate this phenomenon.

We evaluated the efficacy of the phage cocktail delivered by local (i.n.) or systemic (i.p.) routes in a chronic infection model established by direct intratracheal administration of PAO1 with alginate-based beads. Our results showed that i.n. and i.p. administration of phage cocktail increased the animal survival rate (**Figure 4**), reduced the bacterial burden (**Figure 5**) and lung inflammation (**Figure 6 and 7**). These results indicated that both local or systemic phage delivery could control chronic lung infections, suggesting the efficacy and potential of phage therapy as a promising alternative or adjunctive treatment for chronic respiratory infections. These findings are in line with previous animal researches and clinical studies that have demonstrated the safety and efficacy of phage therapy for the treatment of bacterial respiratory infections, including those caused by antibiotic-resistant strains^5, 6, 17, 34, 35^.

Our study also demonstrated that the application of phage cocktail via i.n. or i.p. route in mice is safe. Comparisons of lung tissue histology (**Figure 6**) and proinflammatory cytokine levels (**Figure 7**) in the phage group and the PBS group did not reveal any significant alterations, providing further evidence of the safety of phage therapy administered via local respiratory or systemic routes. These results support the feasibility of using phage therapy as a treatment option for respiratory infections.

While our study demonstrated promising in vivo efficacy of phage therapy, it should be noted that the experimental conditions did not closely resemble the clinical situation due to the short interval between the onset of infection and the start of phage administration. Therefore, the ultimate applicability of this phage cocktail as a treatment for chronic lung infection caused by *P. aeruginosa* still requires further studies from clinical trials.

In conclusion, this study provides evidence that our four-phage cocktail has the potential to be an effective alternative treatment for chronic lung infections caused by MDR strains of *P. aeruginosa*. These findings have significant implications for the future treatment of MDR bacterial infections and suggest that further research into the use of phages as a therapeutic agent is warranted.

## Materials and Methods

### Bacterial strains

*P. aeruginosa* PAO1 strain was kindly provided by Prof. Fan Jin’s group at Shenzhen Institute of Advanced Technology, Chinese Academy of Sciences. Clinical isolates of *P. aeruginosa* were isolated from infected patients at the Third People’s Hospital of Shenzhen, the People’s Hospital of Shenzhen and Southern University of Science and Technology Hospital Shenzhen, China.

### Isolation of Phages

Phages were isolated from various environmental samples by using routine isolation techniques, as previously described^36^. Briefly, bacterial strains were used to isolate and propagate pathogen-specific phages. Following isolation, the phages were triply plaque-purified on their respective host bacterium. Finally, small-scale phage amplification on their corresponding host bacterium was performed to prepare the phage library, which was subsequently stored at 4°C until required.

### Phage preparation

Single colonies of *P. aeruginosa* were picked and the host bacteria were incubated overnight at 37°C. Dilute the overnight culture solution 1:20 and incubate at 37°C for 2 h in a constant temperature shaker at 220 rpm, then add the phage with an MOI 1:100. Add MgCl_2_ and concentrate the phage using a 15 mL 100 kDa ultrafiltration tube, centrifuge 3 000 g, 3-5 min. Add 10 mL SM buffer and centrifuge 3000 g, 3-5 min. Repeatedly blow the concentrate, then aspirate the concentrated phage and store it at 4 °C.

### Phage DNA Extraction, Genome Sequencing, and Assembly

Phage particles were precipitated with 10% polyethylene glycol 8,000 (PEG 8000) at 4°C overnight, centrifuged at 10,000 g for 15 min, and subsequently suspended in SM buffer. Then the concentrated phage particles were treated using DNase I and RNase A (New England BioLab, Massachusetts, USA) to remove bacterial nucleic acids, genomic phage DNA was extracted with MiniBEST Viral RNA/DNA Extraction Kit (Takara, Beijing, China) following manufacturer’s protocol. Whole genome sequencing was performed at the Tianjing Sequencing Center (Novogene, Beijing, China) using the Illumina HiSeq system (Illumina Inc., San Diego, CA, USA). Reads were assembled with SOAPdenovo2^37^. The resulting contigs were uploaded into the RAST server using the RASTtk annotation workflow^38^. Putative functions of the ORFs were further identified with Blast-P based on amino acid sequences. A linear map of the phage was depicted using the Proksee Server (https://proksee.ca/).

### Phage amplification and purification

To amplify phage in a 2 L system, took 50 mL of bacterial solution and added to 2 L LB medium, incubated at 37 ℃, 220 rpm for 2h, then added recovered phage, the ratio of phage to the number of host bacteria is 1:10 and continue to incubate. After complete lysis of the host bacteria, approximately 4 h, the culture was mixed by high-speed centrifugation at 8 000 rpm for 10 min. The supernatant was filtered through a tangential flow system (filter membrane pore size is 0.22 μm). Then, the phage stock solution is concentrated by passing it through a tangential flow system (ultrafiltration membrane pack with a pore size of 100 kDa). After obtaining a sufficient amount of phage concentrate, the phage was further purified by centrifugation through a cesium chloride (CsCl) gradient. A 20 mL phage sample was added to a centrifuge tube with 1.7 g/mL CsCl at the bottom, 1.5 g/mL CsCl in the middle, and 1.3 g/mL CsCl at the top; to construct the concentration gradient, 20 mL of phage concentrate was added first, followed by 4 mL of 1.3 g/mL CsCl solution in a Pasteur pipette, which was slowly injected by touching its head to the bottom of the centrifuge tube, with visible page delamination; followed by 4 mL of 1.5 g/mL CsCl solution at the bottom of the centrifuge tube and finally 4 mL of 1.7 g/mL CsCl solution at the bottom of the centrifuge tube. Centrifuge for 2 h (4 ℃, 24 000 rpm) using a HiMAC ultracentrifuge. After centrifugation, a white-brown layer is visible between 1.3g/mL and 1.5 g/mL. All the white-brown layer was aspirated with a syringe needle about 2 mm below the layer of the phage, approximately 3 mL of liquid can be aspirated from one tube. The phage sample was dialyzed (using a 20 kDa dialysis membrane) using phage preservation buffer (SM buffer) to remove CsCl from the phage sample, using 5 L at a time and changing the dialysate every 4 h at least 3 times to finally obtain high purity phage. The endotoxin concentration of the phage preparations was measured using an endotoxin assay kit (Bioendo, Xiamen, EC80545) following manufacturer’s protocol.

### Preparation of *P. aeruginosa* alginate microbeads

*P. aeruginosa* alginate microbeads were prepared as previously reported with serval modificatcations^31^. PAO1 strain was inoculated onto LB agar medium and incubated in a biochemical incubator for overnight at 37°C. Single colonies were picked from the medium, placed in LB medium, and shaken overnight at 220 rpm at 37°C in a thermostatic shaking incubator. Centrifuged the bacterial culture at 12 000 rpm 4°C for 5 min, added an appropriate amount of LB liquid to the resulting broth, shook and mixed, added 1 mL of PAO1 broth to 8 mL of alginate solution, mixed thoroughly, and poured into an alginate microbead maker, spray it in a mist using compressed air into 0.1 mM Tris-HCl buffer containing 0.1 mol/L CaCl_2_ (pH=7.0) for 1 h, then centrifuged at 700 rpm for 8 min at 4°C, rinsed with sterile 0.9% NaCl containing 0.1 mM CaCl_2_ and repeated twice, and the supernatant discarded to form biofilm-like *P. aeruginosa* microbeads encapsulated by alginate. The suspension was then incubated on agar plates, colonies were counted and the bacterial concentration was adjusted to the desired

1.4*10^8^ CFU/mL and stored at 4°C in the refrigerator.

### Lytic effect of phage against PAO1 in vitro

To determine the bacteriolytic activity of phage, planktonic PAO1 and PAO1 in alginate beads were infected with phage at MOI of 1:100 and 1: 10, respectively. Each group performed in 3 replicates. The plate was incubated at 37℃ continuous oscillation culture and the OD_600_ value was measured every 1h for 24 h in a microplate reader.

### Experimental animals

68 SPF-grade BALB/c mice, male, 6-8 weeks old, weighing 18-22 g, were purchased from Guangdong Vital River Laboratory Animal Technology Co., Ltd. The animal experiments were approved by the Animal Ethics Committee of the Hong Kong Polytechnic University and met the requirements of animal ethics and animal welfare. The mice were randomly grouped in different cages and acclimatized under good routine conditions for 5-6 days before surgery.

### Animal infection model

Mice were fixed on a sterile surgical table, anesthetized with isoflurane, and then a tracheotomy was performed in the middle of the larynx. 50 μL of PAO1 alginate beads suspension was aspirated with a microinjection needle and injected into the lungs through the trachea to infect the lungs. After the injection, the incision was closed using silk sutures. Thirty minutes after the completion of the lung infection model, phage cocktail was given by intranasal (i.n.) and intraperitoneal (i.p.) delivery routes. PBS was given as a control. The phage cocktail was dosed as follows: 5*10^7^ PFU each for Phage PA_ZH1, PA_LZ7, PA_GL1, and PA_CQ9, for a total of 2*10^8^ PFU, MOI=28. The mice were treated with the phage cocktail for 7 days, once daily. Body weight, dietary activity, mental status, and changes in hair were monitored.

### Lung tissue homogenate for bacterial colony counting

After 7 days of *P. aeruginosa* infection, 35 mice in the treatment and control groups were executed by cervical dislocation after anesthesia with isoflurane, the entire thorax and abdomen were routinely disinfected, the skin was cut from under the glabella, the muscle layer was bluntly separated into the abdominal cavity, then the thorax was exposed by cutting along the ribs in an inverted triangle, the lungs were separated, the right lung specimens were collected aseptically, weighed, and the right lung specimens were placed in a homogenized grinding centrifuge tube with 1 mL of pre-cooled PBS solution and prepare lung tissue homogenate at 4°C under ice bath. The tissue homogenate was diluted to 10^−8^ in a 10-fold serial dilution with LB liquid medium. 50 μl of each concentration of homogenate dilution was taken and evenly applied to LB agar solid medium with a smear stick and incubated in a biochemical incubator at 37°C for 24 h for colony counting.

### Histopathological examination of lung

Mice were executed by cervical dislocation method after anesthesia with isoflurane breath, and specimens of the left lung were collected and visualized for gross pathological changes in the lung. The left lung tissue was later fixed in a 4% paraformaldehyde fixative. After dehydration, paraffin embedding, and sectioning for hematoxylin-eosin staining before the histopathological examination, the extent and degree of inflammation were analyzed by light microscopy.

### Lung tissue cytokine assay

After execution of the mice, the right lung specimens were placed in a homogenizing and grinding centrifuge tube under aseptic operation, 1 mL of pre-cooled PBS solution was added and the lung tissue homogenate was prepared at 4°C in an ice bath. After the lung tissue homogenate was obtained, the homogenate was centrifuged at 10 000 rpm for 5 min at a low temperature to obtain the lung tissue homogenate supernatant. The supernatant was used to detect cytokine levels in the lung tissue using the Luminex kit, including TNF-α, IL-6, IL-10, IL-1β, IFN-γ, G-CSF, CCL2/JE/MCP-1 and CCL4/MIP-1β.

### Statistical analysis

Comparisons were performed by Mann-Whitney test except for the survival analysis, where Gehan-Breslow-Wilcoxon test was applied. All statistical analyses were performed using Prism 7.04 (GraphPad, San Diego, CA, USA), and differences with *P*< 0.05 were considered statistically significant.

